# CBD-1 scaffolds two independent complexes required for eggshell vitelline layer formation and egg activation in *C. elegans*

**DOI:** 10.1101/369504

**Authors:** Delfina P. González, Helen V. Lamb, Diana Partida, Zachary T. Wilson, Marie-Claire Harrison, Julián A. Prieto, James J. Moresco, Jolene K. Diedrich, John R. Yates, Sara K. Olson

## Abstract

Metazoan eggs have a specialized coat of extracellular matrix that aids in sperm-egg recognition. The coat is rapidly remodeled after fertilization to prevent polyspermy and establish a more permanent barrier to protect the developing embryo. In nematodes, this coat is called the vitelline layer, which is remodeled into the outermost layer of a rigid and impermeable eggshell. We have identified three key components of the vitelline layer structural scaffold – PERM-2, PERM-4 and CBD-1, the first such proteins to be described in the nematode *C. elegans*. CBD-1 recruited PERM-2 and PERM-4 to the nascent vitelline layer via two N-terminal chitin-binding domains. After fertilization, all three proteins redistributed from the zygote surface to the outer eggshell. Depletion of PERM-2 and PERM-4 from the scaffold led to a porous vitelline layer that permitted soluble factors to leak through the eggshell and resulted in embryonic death. In addition to its role in vitelline layer assembly, CBD-1 is also known to scaffold a protein complex required for fertilization and egg activation (EGG-1-5/CHS-1/MBK-2). We found the PERM complex and EGG complex to be structurally and functionally independent, and regulated through distinct domains of CBD-1. CBD-1 is thus a multifaceted regulator that promotes distinct aspects of vitelline layer assembly and egg activation. In sum, our findings identify the first vitelline layer components in nematodes, and provide a foundation through which to explore both conserved and species-specific strategies used by animals to build protective barriers following fertilization.

## Introduction

Metazoan oocytes are covered by a layer of extracellular matrix that promotes species-specific recognition between egg and sperm. Following fertilization, the matrix is rapidly remodeled in order to prevent polyspermic fertilization and to protect the developing embryo from environmental insults (Wong and Wessel, 2008a). While this protective barrier goes by many different names – the fertilization envelope in sea urchin, the zona in mammals, the chorion in fish, and the eggshell in diptera and nematodes – the remodeling of the matrix utilizes evolutionarily conserved processes, including exocytosis of specialized secretory vesicles called cortical granules; activation of proteases, glycosidases, and cross-linkers; and assembly of carbohydrate- and glycoprotein-based matrices for structural support (Wong and Wessel, 2006).

The nematode eggshell is a multi-layered structure that provides mechanical support and environmental protection to the developing embryo (Christenson, 1950; Johnston and Dennis, 2011; Stein and Golden, 2015). It assembles in a step-wise manner using materials supplied by the oocyte. The outermost vitelline layer is thought to reside on the surface of the unfertilized oocyte, but little is known about its composition other than that it contains glycoproteins (Johnston et al., 2006; Natsuka et al., 2005). Just beneath the vitelline layer, the second layer forms upon fertilization through activation of chitin synthase (CHS-1), which polymerizes and deposits chitin beneath the vitelline layer, likely contributing to lifting of the vitelline layer from the zygote surface (Johnston et al., 2010; Maruyama et al., 2007; Zhang et al., 2005). Soon after, entry into anaphase of meiosis I triggers exocytosis of cortical granules that release their cargo of chondroitin proteoglycans (CPGs) to form the third eggshell layer (Bembenek et al., 2007; Olson et al., 2012; K. Sato et al., 2006; M. Sato et al., 2008). Following anaphase of meiosis II, the innermost layer of the eggshell assembles from fatty acid-based lipids to form the permeability barrier, which limits passage of small molecules and provides osmotic support to the embryo (Benenati et al., 2009; Olson et al., 2012; Rappleye et al., 2003; Tagawa et al., 2001). As one of the most impermeable structures in the animal kingdom (Anya, 1976; Christenson, 1950; Fairbairn, 1957), it is important to understand how the eggshell forms, the degree to which its construction mirrors that of extracellular barriers in other metazoans, and whether it can be exploited to combat parasitic nematode infection.

The outermost vitelline layer of the eggshell sits at an important junction. Proteins embedded in the vitelline layer are likely to play a role in sperm-egg recognition and/or binding, yet after fertilization must be rapidly remodeled in order to prevent polyspermic fertilization and to create the first line of defense in eggshell protection. A protein complex involved in fertilization and polyspermy prevention was recently identified and shown to reside on the oocyte surface. <u>C</u>hitin-<u>b</u>inding <u>d</u>omain protein 1 (CBD-1) anchors the <u>Egg</u> Sterile (Egg) family members EGG-1 and EGG-2 to the cortex, where they have been proposed to play a role in fertilization (Johnston et al., 2010; Kadandale et al., 2005). CBD-1 also helps to anchor CHS-1, the EGG-3/4/5 proteins, and mini-brain kinase (MBK-2) to the oocyte cortex. Soon after fertilization, CHS-1 and the EGG-1/2/3 proteins are endocytosed, while EGG-4/5 and MBK-2 are released from the cortical tether into the cytoplasm to regulate the oocyte-to-embryo transition (Cheng et al., 2009; Maruyama et al., 2007; Stitzel et al., 2007), highlighting the importance of these proteins in egg activation, but not in eggshell vitelline layer remodeling. The fate of CBD-1 after fertilization is unclear; Johnston et al. (Johnston and Dennis, 2011)proposed that chitin polymer secreted at fertilization could bind the chitin-binding domains of CBD-1 and relieve its tether to the membrane-bound EGG-1-5/CHS-1/MBK-2 complex (hereafter called the EGG complex), but experiments to support this model have not yet been conducted. If CBD-1 is released from the zygote surface after fertilization, it is likely that proteins other than members of the EGG complex assist CBD-1 to remodel the nascent vitelline layer into the outermost layer of the eggshell.

To investigate vitelline layer formation in *C. elegans*, we examined proteins from a previous RNAi screen that were promising candidates for vitelline layer formation (Carvalho et al., 2011). We found that <u>Perm</u>eable Eggshell protein 2 (PERM-2), PERM-4, and CBD-1 are essential components of the eggshell vitelline layer. Recruitment of PERM-2 and PERM-4 to the vitelline layer requires two chitin-binding domains in the N-terminus of CBD-1, but these two domains and the PERM complex itself are dispensable for the EGG complex, highlighting the dual role of CBD-1 in vitelline layer formation and egg activation. Disruption of PERM-2 and PERM-4 function by RNAi or null mutation rendered the eggshell permeable to dyes and caused embryonic lethality, indicating that PERM-2 and PERM-4 maintain the structural integrity of the vitelline layer that is important for establishing the impermeability of the nematode eggshell. In sum, this study characterizes the first set of bona fide vitelline layer proteins in *C. elegans* and lays the groundwork for future studies that compare vitelline layer remodeling in nematodes to other metazoans.

## Materials & Methods

### *C. elegans* strains

Worm strains used in this study are listed in Table S1, some of which were provided by the CGC, funded by NIH Office of Research Infrastructure Programs (P40 OD010440). N2 (wild-type) (Brenner, 1974), AD189 (GFP::EGG-1) and RT495 (EGG-2::GFP) (Kadandale et al., 2005), AD200 (EGG-3::GFP) and AD238 (EGG-3::mCherry) (Maruyama et al., 2007), OD95 (GFP::PH^PLC*1δ1*^; mCherry:: H2B) (Audhya et al., 2005), OD344 (mCherry::CPG-2) and OD367 (mCherry::CPG-1; GFP:: PH) (Olson et al., 2012), and VC2258 (*cbd-1(ok2913)* mutant) (C. elegans Deletion Mutant Consortium, 2012) were previously described. AG212 (CBD-1::mCherry) was a generous gift from David Levine and Andy Golden (NIH NIDDK). Strains were maintained at 20°C-25°C on nematode growth media (RPI, N81800) seeded with OP50 bacteria.

### Plasmids and transgenic strains

PERM-2::mCherry (POM1) and PERM-4::mCherry (POM6) strains were generated via Mos1-mediated single copy insertion (MosSCI) as described (Frøkjær-Jensen et al., 2008). Gibson assembly (Gibson et al., 2009)was used to combine PCR products amplified by Q5 DNA Polymerase (NEB, M0491) with the primers in Table S2 using pBluescript, N2 genomic DNA, or pAA65 (McNally et al., 2006) as DNA sources for the vector backbone, *perm-*2 or *perm-4*, and mCherry sequences, respectively. AvrII (NEB, R0174) sites were introduced to facilitate subcloning from pBluescript into pCFJ151 that targets a Mos1 transposon site in chromosome II (Frøkjær-Jensen et al., 2008). pSO53 contained *perm-2* coding sequence fused to a C-terminal mCherry tag under the control of its endogenous regulatory sequences, while pSO58 contained similar sequences for C-terminally tagged *perm-4*. Paralyzed EG6429 adult worms (*unc-119(ed3)*; *ttTi5605 II*) (Frøkjær-Jensen et al., 2014) were injected in the gonad with a mixture containing pCFJ601 (Mos1 transposase); pMA122 (*peel-1*) (Seidel et al., 2011); co-injection markers pCFJ90, pCFJ104, and pGH8; and either pSO53 or pSO58 (Frøkjær-Jensen et al., 2014; 2008). One week after injection the animals were heat shocked at 34°C for 2hr to activate *peel-1* counter-selection, and moving worms were screened for absence of co-injection markers. Properly integrated lines were verified by DNA sequencing and outcrossed six times to N2 males.

The *perm-2(pmn1)* null allele (POM37) was created via CRISPR/Cas9-mediated gene editing according to the long-range homology-directed repair and self-excising cassette (SEC) method (Dickinson et al., 2015) using the oligos listed in Table S2. The pSO210 sgRNA/Cas9 plasmid to introduce a double-stranded break near the start codon of *perm-2* was generated by site-directed mutagenesis of plasmid pJW1219 (Ward, 2015). The pSO212 repair template was designed to insert GFP in place of the *perm-2* coding sequence, and was created by cutting pDD282 with ClaI/SpeI and performing HiFi DNA assembly (NEB, E5520) with PCR-generated left (557 bp) and right (608 bp) homology arms containing 30 bp of overlap with the pDD282 vector (Dickinson et al., 2015). Adult N2 worms were injected with a mixture containing pSO210, pSO212, and co-injection markers pCFJ90, pCFJ104, and pGH8. Hygromycin (5 mg/ml, Thermo Fisher, 10-687-010) was added to growth plates on day 3 post-injection to select for animals harboring the SEC, and one week later animals were screened for genomic edits through presence of the SEC (Rol phenotype) and absence of co-injection markers. Three independent lines were obtained, and one was selected for six outcrosses to N2 and SEC excision. Correct removal of *perm-2* coding sequence and replacement with GFP was verified by sequencing through the edited region (Eurofins-Operon). All vectors for MosSCI and CRISPR/Cas9 cloning were obtained from Addgene.

### RNA interference

Double-stranded RNA was prepared by PCR amplification of N2 genomic DNA using the oligos listed in Table S2, and *in vitro* transcription with Megascript T7 and T3 kits (Thermo Fisher, AM1334 and AM1338). RNA was purified by phenol-chloroform extraction and isopropanol precipitation. Pellets were solubilized in 1x soaking buffer (11mM Na_2_HPO_4_, 5.5mM KH_2_PO_4_, 2.1mM NaCl, 4.7mM NH_4_Cl) and annealed by incubation at 68°C for 10 minutes followed by 37°C for 30 minutes. Animals at the L4 or young adult stage were injected with 1 mg/ml dsRNA and grown for 6-30 hours at 20-23°C for imaging experiments, or recovered for 2 hours at room temperature and singled to individual plates for brood size and permeability assays. Control RNAi experiments were performed with 1 mg/ml of *him-8* dsRNA or 0.5 mg/ml each of dsRNA targeting *cpg-1* and *cpg-2*; neither control affects vitelline layer formation (Olson et al., 2012) (Olson et al., 2006).

### Brood size and eggshell permeability assays

Injected L4-stage worms were singled onto individual plates and allowed to lay eggs for three days with transfer to a fresh plate each day. For permeability assays, growth plates were supplemented with 150 ug/ml Nile Blue A (Sigma, N0766). Twenty-four hours after removal of the adult worm, plates were scored for number of hatched larvae and unhatched dead eggs (+/-blue dye for permeability assays). Brood size was defined as the sum of hatched larvae over three days of laying. Permeability was defined as (blue embryos)/(larvae + clear dead embryos + blue dead embryos) (Carvalho et al., 2011).

### Immunofluorescence

For immunofluorescence experiments, embryos were dissected on subbing solution-coated slides (4 mg/ml gelatin USP, 0.4 mg/ml chromalum, 1 mg/ml poly-L-lysine) in 0.75x egg salts (1x egg salts: 118 mM NaCl, 40 mM KCl, 3.4 mM MgCl_2_, 3.4 mM CaCl_2_, 5 mM Hepes pH7.4), freeze cracked, and fixed in −20°C methanol for 15 minutes as described previously (Olson et al., 2012). Following rehydration in PBS, slides were stained overnight with a 1:100 dilution of rhodamine-conjugated chitin-binding probe (New England Biolabs, discontinued) and 1 ug/ml DAPI (ThermoFisher, D1306) before being mounted in 0.5% p-phenylenediamine in 90% glycerol, 20 mM Tris pH 8.8. In some cases, the eggshell vitelline layer was removed prior to the freeze-cracking step by brief alkaline-bleach treatment (0.5 N NaOH, 2.5% bleach in M9 buffer). In all cases, a single central plane was imaged.

### Live imaging microscopy

Whole worm imaging was performed by anesthetizing adults in a 2 ul drop containing 3 mg/ml tricane and 0.3 mg/ml tetramasole in M9 buffer on a multiwell test slide (MP Biomedicals, 096041505). Embryo imaging was performed by dissecting adults on a 24×50mm cover glass in a 3 ul drop of 0.75x egg salts surrounded by a thin ring of petroleum jelly, which prevented compression when overlayed with 22×22 mm cover glass (Olson et al., 2012). In some experiments egg salts were supplemented with 6 uM FM4-64 or 2 ug/ml DAPI to verify eggshell permeability. Images of a single central plane were collected on an upright Nikon Eclipse 90i microscope equipped with a QuantEM 512SC CCD camera (Photometrics) using a 40×0.75 NA Plan Fluor or 100×1.40 NA Plan Apo VC objective lens and analyzed with either Nikon Elements or Fiji software (Schindelin et al., 2012).

Variance values in Fig. 3C-E were measured by drawing a line along the boundary between two adjacent oocytes and obtaining plot profiles in Nikon Elements. The variance in log_10_ fluorescence intensity ratios between mCherry and GFP signals along the line was calculated to determine whether the mCherry-tagged vitelline layer protein exhibited clustering patterns similar to or different from the GFP::PH plasma membrane signal.

### Co-immunoprecipitation and protein identification by mass spectrometry

Crude extract was prepared from POM24 (PERM-4::mCherry; GFP::PH) and POM27 (PERM-2::mCherry; GFP::PH) adult worms by sonication in 10 mM Tris pH 7.5, 150 mM NaCl, 0.5 mM EDTA and 0.5% NP-40, and the soluble fraction was incubated with RFP-Trap magnetic agarose beads as recommended by the manufacturer (Chromotek). Immunoprecipitated proteins were eluted in 50 mM Tris pH8.5 containing 8M urea (Cheeseman et al., 2004) and processed for protein identification by mass spectrometry (Fonslow, 2014).

For mass spectrometry analysis, samples were precipitated by methanol/chloroform. Dried pellets were dissolved in 8 M urea/100 mM TEAB, pH 8.5. Proteins were reduced with 5 mM tris(2-carboxyethyl)phosphine hydrochloride (TCEP, Sigma-Aldrich) and alkylated with 10 mM chloroacetamide (Sigma-Aldrich). Proteins were digested overnight at 37°C in 2 M urea/100 mM TEAB, pH 8.5, with trypsin (Promega). Digestion was quenched with formic acid, 5% final concentration. The digested samples were analyzed on a Q Exactive mass spectrometer (Thermo). The digest was injected directly onto a 15cm, 100um ID column packed with Aqua 3um C18 resin (Phenomenex). Samples were separated at a flow rate of 300nl/min on an Easy nLCII (Thermo). Buffer A and B were 0.1% formic acid in 5% acetonitrile and 0.1% formic acid in 80% acetonitrile, respectively. A gradient of 1-35% B over 180 min, an increase to 80% B over 40 min, and held at 90% B for 5 min of washing prior to returning to 1% B was used for 240 min total run time. The column was re-equilibrated with 10ul of buffer A prior to the injection of sample. Peptides were eluted directly from the tip of the column and nanosprayed directly into the mass spectrometer by application of 2.5kV voltage at the back of the column. The Q Exactive was operated in a data dependent mode. Full MS^1^ scans were collected in the Orbitrap at 70K resolution with a mass range of 400 to 1800 m/z. The 10 most abundant ions per cycle were selected for MS/MS and dynamic exclusion was used with exclusion duration of 15 sec.

Protein and peptide identification were done with Integrated Proteomics Pipeline – IP2 (Integrated Proteomics Applications). Tandem mass spectra were extracted from raw files using RawConverter (He et al., 2015) and searched with ProLuCID (Xu et al., 2015) against a protein database from Wormbase.org (PRJNA13758.WS265) appended with common contaminants and RFP fusion proteins. The search space included all fully-tryptic and half-tryptic peptide candidates. Carbamidomethylation on cysteine was considered as a static modification. Data was searched with 50 ppm precursor ion tolerance and 600 ppm fragment ion tolerance. Identified proteins were filtered using DTASelect (Tabb et al., 2002) and utilizing a target-decoy database search strategy to control the false discovery rate to 1% at the protein level.

### Statistics and protein predictions

Statistical analysis was carried out using unpaired student’s t-tests with unequal variance using GraphPad software (www.graphpad.com/quickcalcs/ttest1.cfm). O-linked mucin attachment sites were predicted based on consensus sequence (Chen et al., 2008). Chitin-binding domains were predicted by Prosite (prosite.expasy.org).

Critical reagents and resources are compiled in the Key Resources Table.

## Results

### PERM-2 and PERM-4 are required for eggshell integrity

PERM-2 and PERM-4 were previously identified in an RNAi screen for proteins required for eggshell formation in *C. elegans* (Carvalho et al., 2011). PERM-2 and PERM-4 are conserved in free-living (*Caenorhabditis*, *Pristionchus*) and parasitic nematode genera (*Strongyloides*, *Onchocerca*), but little is known about their structure or function. PERM-2 and PERM-4 contain signal sequences and predicted O-linked mucin glycan attachment sites (Chen et al., 2008), so are likely to be secreted, but lack any other conserved domains (Fig. 1A). To further investigate their function in eggshell formation, we analyzed the effect of PERM-2 and PERM-4 depletion on brood size and eggshell permeability. Inhibition of PERM-2 or PERM-4 by RNAi or null mutation significantly decreased brood size and increased eggshell permeability (Fig. 1B), indicating these proteins play an important role in maintaining the integrity of the *C. elegans* embryonic eggshell. Given that the *perm-2(null)* mutation had a less severe phenotype than RNAi inhibition, we decided to investigate onset of penetrance by analyzing brood size and permeability over the three successive days of adult fertility. Eggshell permeability defects became increasingly severe over time, particularly between days 1 and 2 of adulthood (Fig. 1C), suggesting that a factor able to compensate for the function of PERM-2 dropped below a threshold level. Embryos depleted of PERM-2 and PERM-4 readily incorporated FM4-64 dye from the surrounding media into their plasma membranes, indicating that the eggshell permeability barrier was compromised (Fig. 1D). Simultaneous depletion of both PERM-2 and PERM-4 failed to enhance the phenotype and embryos were able to complete several divisions before arresting around the gastrulation stage (Fig. 1D), a phenotype associated with inhibition of proteins that form the innermost permeability barrier of the eggshell (Olson et al., 2012).

**Figure 1.**
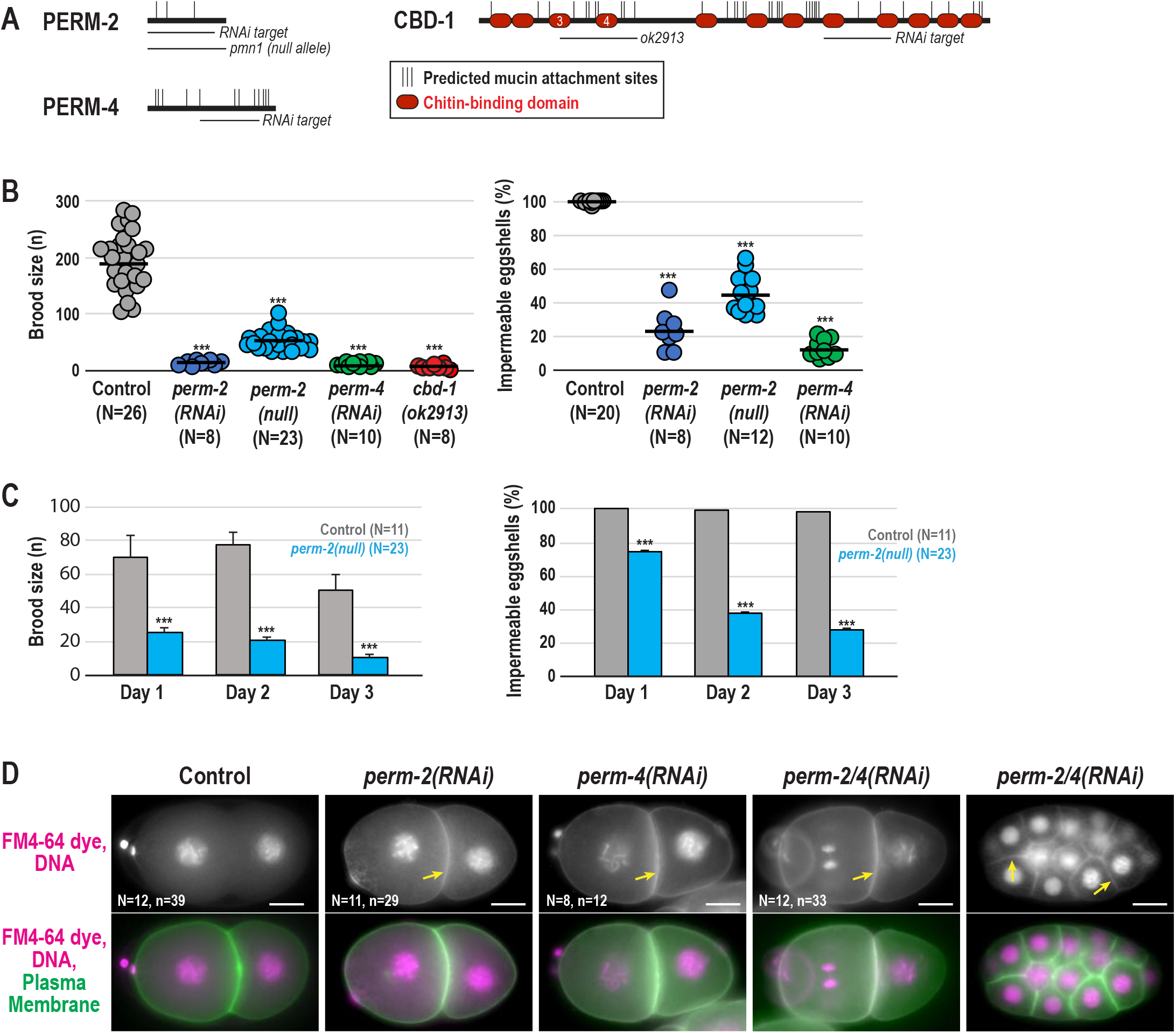
PERM-2 and PERM-4 are required for eggshell integrity. **(A)** Schematic representations of PERM-2, PERM-4 and CBD-1 showing the location of predicted O-linked mucin attachment sites (thin vertical black lines), chitin-binding domains (red ovals), deletion allele, and target regions for dsRNA-mediated interference (RNAi). Chitin-binding domains 3 and 4 of CBD-1 are disrupted by the in-frame *ok2913* deletion allele. **(B)** Left, brood size counts calculated as the number of hatched progeny from a single worm over a three-day laying period. Right, eggshell impermeability represents the fraction of embryos laid by a single adult that were able to exclude Nile Blue dye. **(C)** Progressive penetrance in the *perm-2(null)* strain (blue bars) of defects in average brood size (left) and ability to produce impermeable eggshells (right) for three successive days after initial plating of L4-stage worms, compared to controls (gray bars). For all graphs, data were pooled from 2-3 independent experiments per condition. Control worms were either N2 wild-type worms (deletion allele experiments) or injected with *him-8* dsRNA (RNAi experiments); control data were pooled since brood size means and standard errors were similar (198±16 N2 N=11 vs. 188±13 *him-8 dsRNA* N=15). N=number of adults, n=number of hatched larvae. Horizontal black bars in (B), means. Error bars in (C), standard error of the mean (SEM). ***, p<0.0001 compared to controls by unpaired student’s t-test with unequal variance. **(D)** Widefield fluorescence images of control and RNAi-treated embryos expressing mCherry-tagged histone H2B (magenta) and a GFP-tagged plasma membrane marker (green) that were dissected into FM4-64 dye (magenta). Arrows highlight plasma membrane stained with FM4-64 dye in permeable embryos. Multicellular embryo shown for *perm-2/4(RNAi)* highlights its ability to complete multiple divisions before arresting. N=number of dissected adults, n=number of embryos. Scale bars, 10um.

### PERM-2 and PERM-4 localization to the vitelline layer is co-dependent and requires CBD-1

Given that PERM-2 and PERM-4 inhibition resulted in a permeability barrier-like defect, we sought to determine whether these proteins localized to the permeability barrier. Surprisingly, mCherry-tagged PERM-2 and PERM-4 were found on the surface of oocytes in the gonad before fertilization and the embryonic eggshell after fertilization (Fig. 2AB), a pattern consistent with the outermost vitelline layer of the eggshell rather than the innermost permeability barrier (Johnston and Dennis, 2011; Olson et al., 2012; Stein and Golden, 2015). The chitin-binding domain protein CBD-1 similarly localized to unfertilized oocytes in a previous study (Johnston et al., 2010) and is proposed to be a vitelline layer protein (Johnston and Dennis, 2011; Stein and Golden, 2015), so we also investigated its localization following fertilization. Similar to PERM-2 and PERM-4, CBD-1::mCherry also redistributed from the oocyte membrane to the embryonic eggshell after fertilization (Fig. 2C). In order to verify vitelline layer localization, we treated PERM-2::mCherry, PERM-4::mCherry and CBD-1::mCherry embryos with alkaline bleach, which is known to remove the outer vitelline layer from the eggshell (Rappleye et al., 1999). All three markers were successfully stripped from the eggshell following bleach treatment (Fig. 2ABC). We have thus identified PERM-2, PERM-4, and CBD-1 as the first bona fide vitelline layer proteins in the *C. elegans* eggshell.

**Figure 2.**
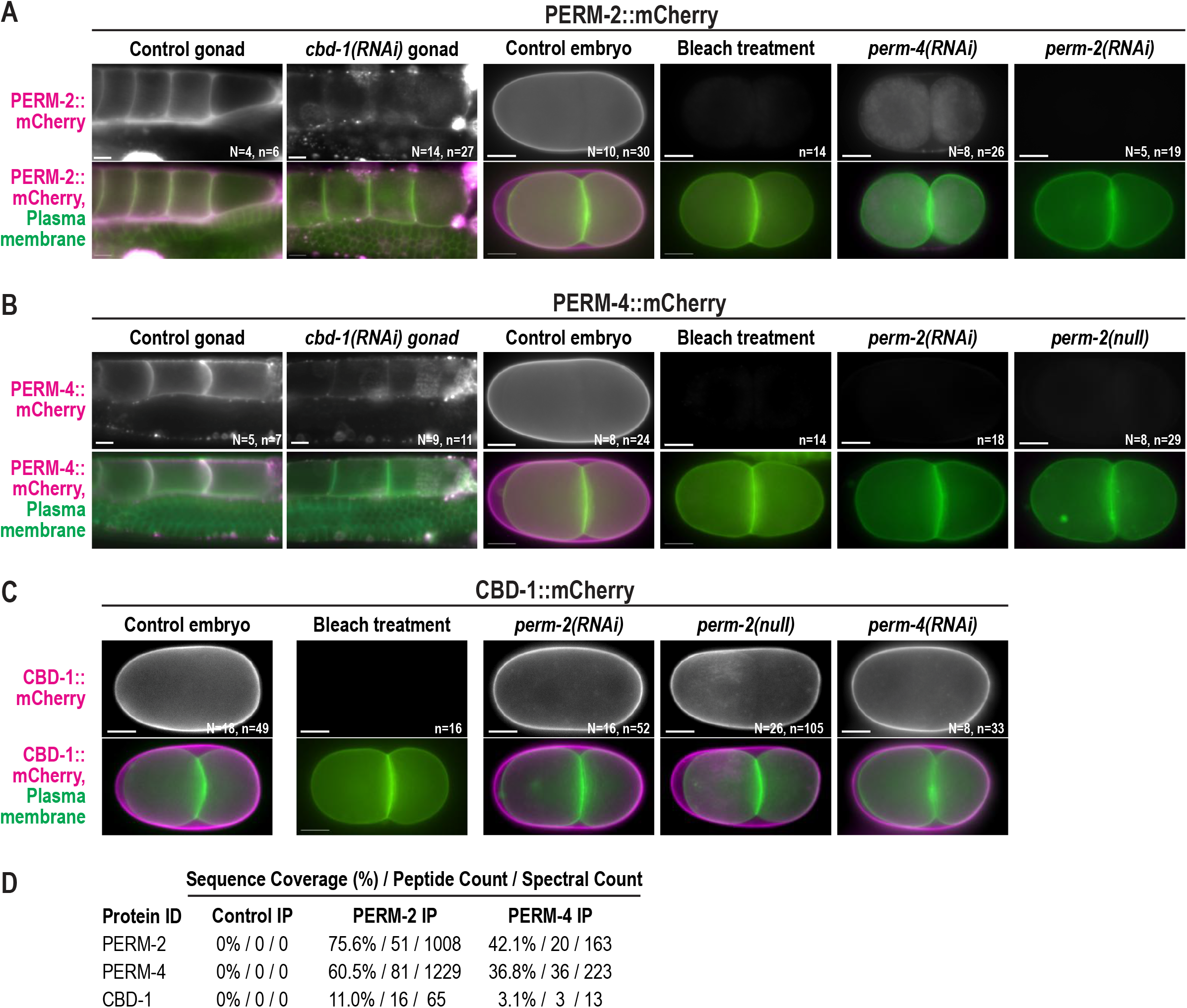
PERM-2 and PERM-4 localize co-dependently to the eggshell vitelline layer and require CBD-1. Widefield fluorescence images of gonads and embryos from control, RNAi-treated, or *perm-2(null)* worms that express a GFP-tagged plasma membrane probe (green) with either PERM-2::mCherry **(A)**, PERM-4::mCherry **(B)**, or CBD-1::mCherry **(C)** (magenta). Some worms were treated with alkaline bleach, which removes the outermost vitelline layer but not the chitin or CPG layers of the eggshell. N=number of adults (not determined for some experiments), n=number of gonads or embryos. Scale bars, 10um. **(D)** Mass spectrometry analysis of proteins co-immunoprecipitated from whole worm extract. Control IP, non-specific IgG beads incubated with PERM-2:mCherry extract. PERM-2 IP, RFP-Trap beads incubated with PERM-2::mCherry extract. PERM-4 IP, RFP-Trap beads incubated with PERM-4::mCherry extract. Table shows sequence coverage, peptide counts, and spectral counts of PERM-2 (or PERM-2::mCherry), PERM-4 (or PERM-4::mCherry), and CBD-1 in the eluted fractions.

Due to their similar localization patterns, we hypothesized that PERM-2, PERM-4 and CBD-1 were part of an interdependent protein complex that provides the structural basis of the vitelline layer. In support of this idea, depletion of PERM-4 disrupted PERM-2::mCherry localization to the vitelline layer, with redistribution of some of the PERM-2::mCherry signal into intracellular compartments (Fig. 2A). Loss of PERM-2 via RNAi or null mutation likewise prevented vitelline layer targeting of PERM-4::mCherry (Fig. 2B), which was subsequently released into the gonad and uterine cavities (data not shown). These data show that PERM-2 and PERM-4 are co-dependent for recruitment and/or maintenance on the vitelline layer. Similarly, depletion of CBD-1 resulted in loss of PERM-2::mCherry and PERM-4::mCherry from the oocyte surface (Fig. 2AB). However, inhibition of PERM-2 and PERM-4 had no effect on CBD-1::mCherry localization (Fig.2C). To explore whether the genetic interactions among PERM-2, PERM-4 and CBD-1 were due to a physical association, we performed co-immunoprecipitation (co-IP) experiments with RFP-Trap followed by protein identification by mass spectrometry. Co-IP of PERM-2::mCherry pulled down PERM-4 and CBD-1, while co-IP of PERM-4::mCherry pulled down PERM-2 and CBD-1. Based on the combination of localization and interaction data, we conclude that CBD-1 functions at the top of the structural hierarchy to recruit PERM-2 and/or PERM-4 to the vitelline layer, and that PERM-2 and PERM-4 cooperate to maintain each other’s association with CBD-1 through physical association between all members of the protein complex

### The PERM complex and EGG complex are structurally independent

A previous study identified CBD-1 as an essential component of a protein complex on the oocyte membrane that organizes factors involved in fertilization and egg activation (Johnston et al., 2010). CBD-1 anchors EGG-1, EGG-2, and CHS-1 to the oocyte membrane and evenly distributes the downstream components EGG-3, EGG-4, EGG-5, and MBK-2 along the cortex (hereafter called the EGG complex) (Cheng et al., 2009; Parry et al., 2009). Given the role of CBD-1 in also recruiting PERM-2 and PERM-4 (hereafter called the PERM complex) to the oocyte surface, we sought to determine whether the EGG complex and PERM complex localize independently to CBD-1, or whether they are structurally interdependent. Inhibition of PERM-2 or PERM-4 had no effect on membrane localization of EGG-1::GFP (Fig. 3A) or cortical localization of EGG-3::GFP (Fig. 3B) in unfertilized oocytes. RNAi co-depletion of EGG-1 and EGG-2 likewise had no effect on the recruitment of PERM-4::mCherry to the oocyte surface (Fig. 3C). The distribution of PERM-2::mCherry showed some accumulation of irregular patches of signal between oocyte boundaries in the absence of EGG-1/ 2 (Fig. 3D). However, CBD-1::mCherry distribution was similarly affected (Fig. 3E), suggesting that EGG-1/2 depletion disrupted CBD-1::mCherry localization, which had an indirect effect on PERM-2::mCherry organization. These data suggest the PERM complex and EGG complex are structurally independent despite both complexes being scaffolded by a common protein, CBD-1.

**Figure 3.**
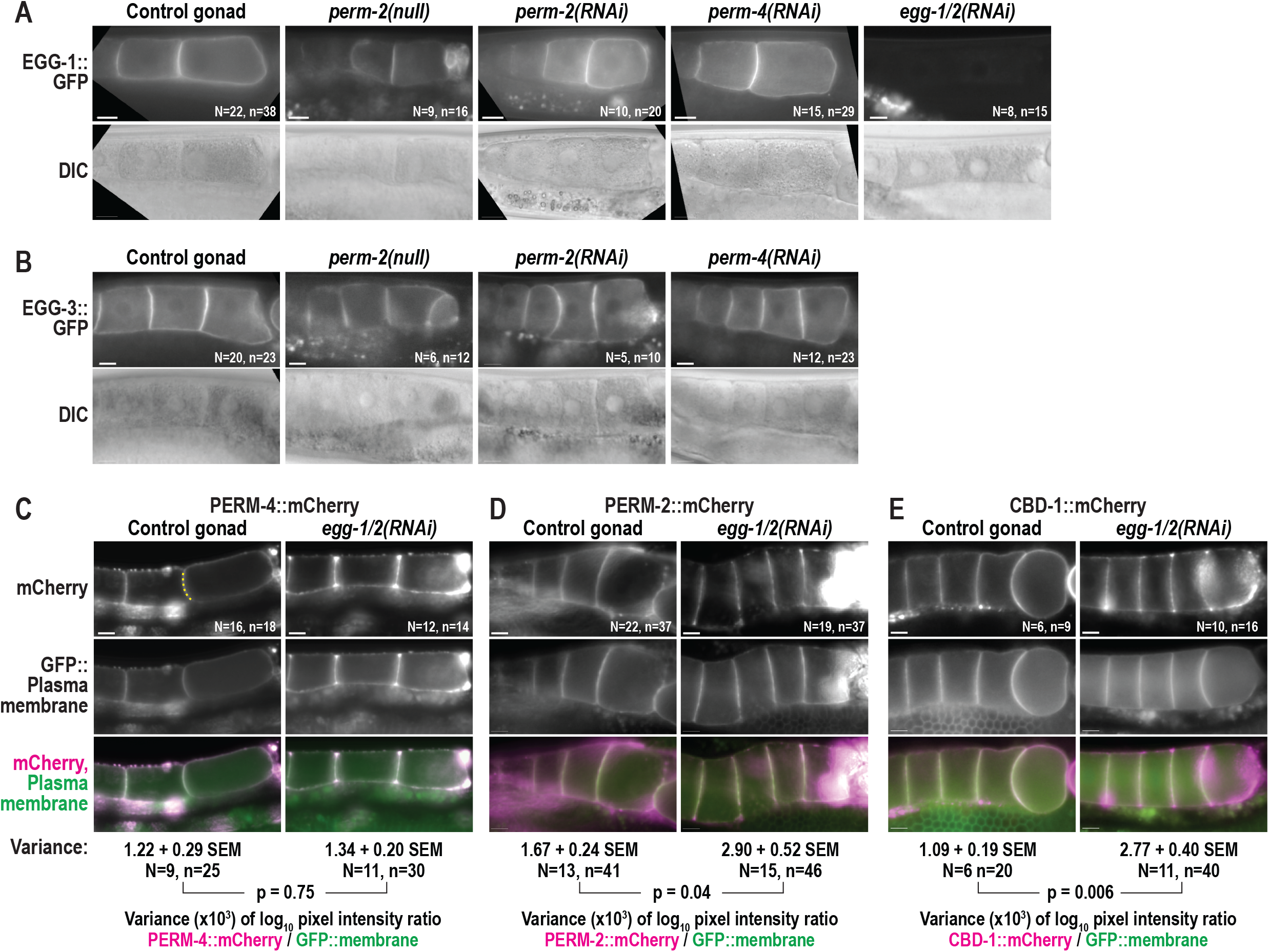
The PERM complex and EGG complex are structurally independent. Widefield fluorescence and DIC images of gonads from control, RNAi-treated, or *perm-2(null)* worms that express EGG-1::GFP **(A)**, EGG-3::GFP **(B)**, PERM-4::mCherry **(C)**, PERM-2::mCherry **(D)**, or CBD-1::mCherry **(E)**. N=number of adults, n=number of gonads. For **(C)-(E)**, irregular distribution of fluorescent markers was calculated as variance in log_10_ fluorescence intensity ratios between mCherry-tagged proteins and a GFP::plasma membrane marker along oocyte boundaries (dotted yellow line in **C**). SEM, standard error of the mean. N=number of gonads, n=number of oocyte boundaries. P-values calculated by unpaired student’s t-test with unequal variance. Scale bars, 10um.

### The PERM complex is scaffolded by the N-terminus of CBD-1

Because our data showed that CBD-1 organizes two independent protein complexes at the oocyte surface, we next hypothesized that distinct regions of the CBD-1 protein may separately scaffold the PERM and EGG complexes. CBD-1 is a large protein (1319 aa, 144 kD) comprised of 12 chitin-binding domains interspersed by consensus sites for O-linked mucin chains (Fig. 1A) (Chen et al., 2008). The *ok2913* deletion allele maintains the *cbd-1* reading frame but disrupts chitin-binding domains #3 and #4 in the N-terminal half of the protein (C. elegans Deletion Mutant Consortium, 2012), allowing us to probe the effect of these chitin-binding domains on recruitment of the PERM and EGG complexes. When the entire CBD-1 protein was depleted by RNAi, EGG-1::GFP redistributed from the oocyte membrane to intracellular vesicles that accumulated in the cytoplasm and near the nuclear envelope (Fig. 4A), and EGG-3::GFP showed irregular distribution along the cortex rather than a continuous layer (Fig. 4B), as previously described (Johnston et al., 2010). However, the *cbd-1(ok2913)* mutant was able to effectively recruit EGG-1::GFP and EGG-3::mCherry to the oocyte membrane, maintained EGG-1::GFP localization during fertilization, and internalized EGG-3::mCherry with normal dynamics in recently fertilized embryos (Fig. 4AB). By contrast, *cbd-1(ok2913)* mutants showed a dramatic decrease in recruitment of PERM-2::mCherry and PERM-4::mCherry to the oocyte surface and the vitelline layer of the eggshell (Fig. 4CD), similar to the effect seen in *cbd-1(RNAi)* animals (Fig. 2AB). In some instances, there was complete failure to recruit PERM-2::mCherry or PERM-4::mCherry (embryo in Fig. 4C and gonad in Fig. 4D), while in other cases there was reduced recruitment of the marker (embryo in Fig. 4D) or redistribution from the membrane surface to soluble regions where oocyte membranes failed to adhere (gonad in Fig. 4C). Inhibition of PERM-2/4 recruitment became stronger as the mutant animals aged, which is consistent with the ability of *cbd-1(ok2913)* worms to generate a handful of viable embryos before the onset of significant embryonic lethality (Fig. 1B). These results suggest that the N-terminus of CBD-1 helps to recruit the PERM complex, but not the EGG complex, to the oocyte before fertilization.

**Figure 4.**
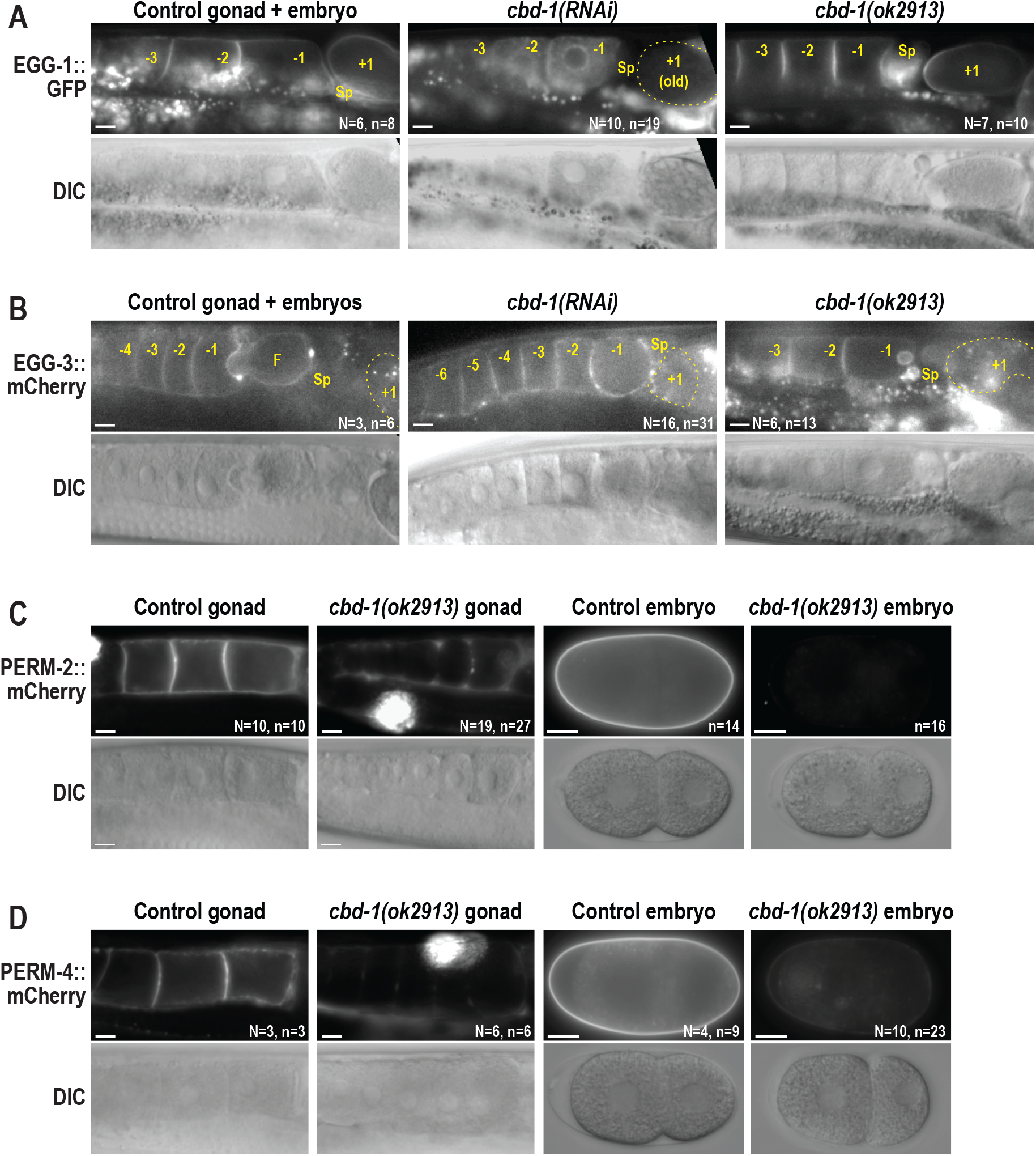
The N-terminus of CBD-1 scaffolds the PERM complex, but not the EGG complex. Widefield fluorescence and DIC images of gonads and embryos from control, RNAi-treated, or *cbd-1(ok2913)* worms that express EGG-1::GFP **(A)**, EGG-3::mCherry **(B)**, PERM-2::mCherry **(C)**, or PERM-4::mCherry **(D)**. Small bright puncta in (A) and (B) represent autofluorescence of gut granules distinct from the EGG-1::GFP or EGG-3::GFP signal in the gonad. Oocytes and embryos are labeled in reference to distance from the spermatheca (Sp), with the −1 oocyte the next to be fertilized, and the +1 embryo (in some cases outlined in a dotted yellow line) recently fertilized and developing in the uterus. Large, bright signal in *cbd-1(ok2913)* mutant gonads in (C) and (D) is accumulation of secreted mCherry fusion protein in coelomocytes. The *cbd-1(ok2913)* allele disrupts the third and fourth chitin-binding domains of CBD-1 (see Fig. 1A). N=number of adults, n=number of gonads or embryos. Scale bars, 10um.

### PERM-2 and PERM-4 promote structural integrity of the vitelline layer

Our existing model for *C. elegans* eggshell assembly states that the eggshell forms in a hierarchical manner, where external layers are essential for assembly of more internal layers (Olson et al., 2012). This model predicts that defects in the outermost vitelline layer would be catastrophic for eggshell assembly, as is observed in *cbd-1(RNAi)* embryos (Johnston et al., 2010). To test this prediction, we examined the structure and function of various markers for other eggshell layers following inhibition of PERM-2 and PERM-4. The chitin layer normally forms immediately beneath the vitelline layer and can be visualized with the external application of a chitin-binding probe (Zhang et al., 2005). Interestingly, depletion of PERM-2 or PERM-4 by RNAi resulted in enhanced staining of the chitin layer, similar to when the vitelline layer is removed by alkaline bleach treatment of wild-type embryos (Fig. 5A) (Zhang et al., 2005). The staining was intense enough that identical exposures of unbleached control embryos failed to detect chitin staining. These results suggest that the chitin layer can form in the absence of PERM-2 and PERM-4, but the chitin probe has greater access to underlying eggshell layers, likely due to an increase in permeability of the vitelline layer. Further evidence of abnormal vitelline layer formation was seen when embryos dissected from PERM-2 or PERM-4 depleted animals extruded from the uterus in long, interconnected chains (Fig. 5B), which is not observed in control animals.

**Figure 5.**
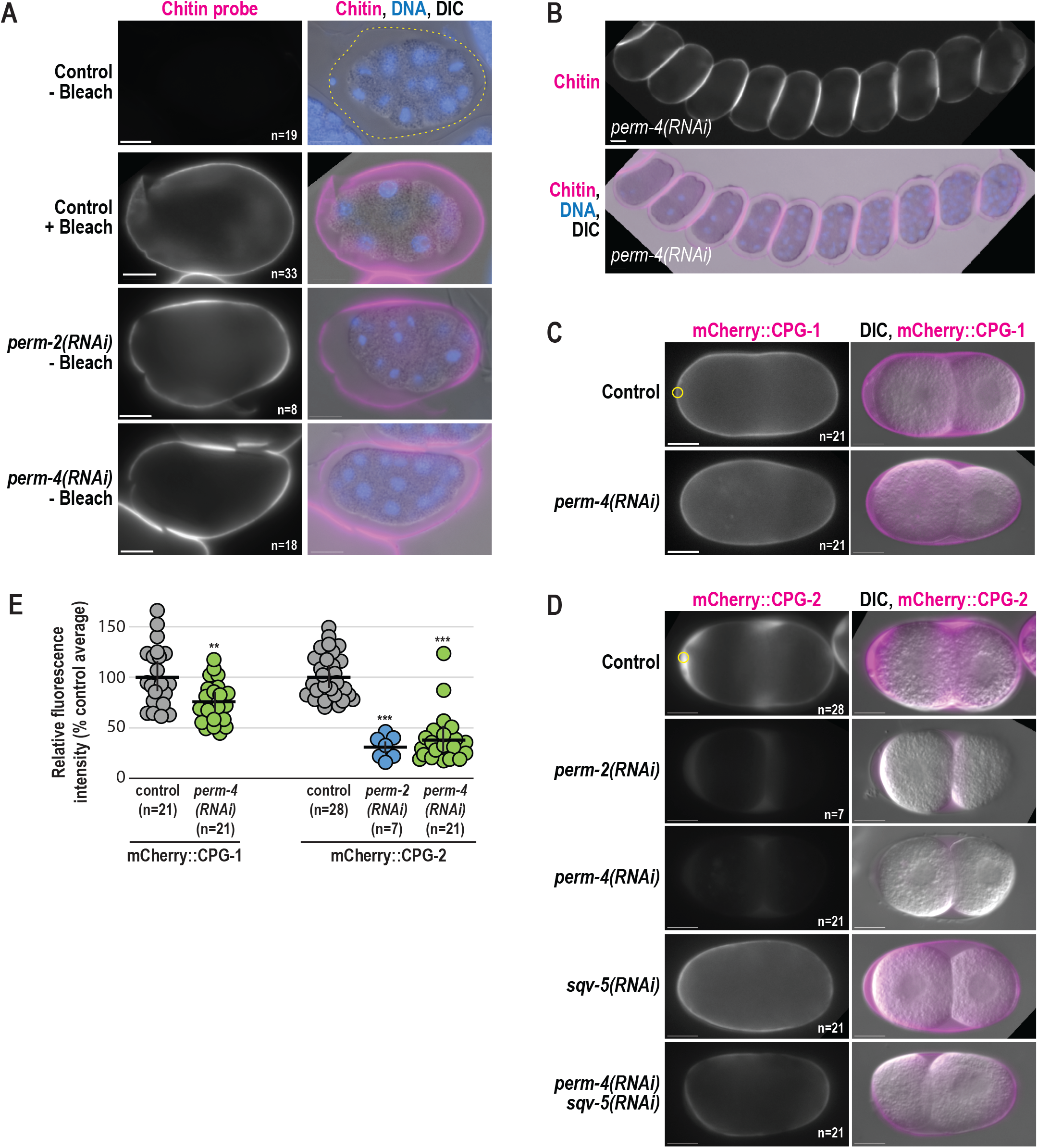
PERM-2 and PERM-4 promote structural integrity of the vitelline layer. **(A,B)** Immunofluorescence of fixed embryos stained with a rhodamine-conjugated chitin-binding probe to mark the eggshell (magenta) and DAPI to mark DNA (blue). **(C,D)** Widefield fluorescence and DIC images of live embryos from control and RNAi-treated worms that express mCherry::CPG-1 **(C)** or mCherry::CPG-2 **(D)**. n=number of embryos. Scale bar, 10um. **(E)** Quantification of relative fluorescence intensity of eggshell markers within a region of interest (ROI, yellow circles) normalized to the control average. Horizontal black bars represent the mean, vertical black bars the standard error of the mean (SEM). **, p=0.0033; ***, p<0.0001 compared to controls by unpaired student’s t-test with unequal variance. n=number of embryos. Scale bars, 10um.

Given the increased permeability to externally-applied factors like the chitin probe, we next explored whether factors secreted from the embryo would be retained by a compromised vitelline layer. Soon after the chitin layer forms, cortical granule exocytosis releases the chondroitin proteoglycans CPG-1 and CPG-2. CPG-1 assembles beneath the chitin layer to form the third layer of the eggshell, while the majority of CPG-2 remains soluble and hydrates the perivitelline space between the eggshell and embryo proper (Olson et al., 2012). In embryos depleted of PERM-2 or PERM-4, mCherry::CPG-1 properly incorporated into the eggshell but at slightly lower levels (Fig. 5CE). By contrast, mCherry::CPG-2 showed a significantly larger decrease in signal (Fig. 5DE), suggesting either that CPG-2 was not secreted into the perivitelline space in the absence of PERM-2, or that the protein failed to be retained in the perivitelline space and leaked out of the eggshell. To discriminate between these hypotheses, we depleted *<u>Sq</u>*uashed *<u>V</u>*ulva 5 (SQV-5), the enzyme required to build chondroitin chains on the CPG core proteins, rendering mCherry::CPG-2 in an unglycosylated and less soluble state (Olson et al., 2012). When SQV-5 and PERM-4 were co-depleted by RNAi, the insoluble form of mCherry::CPG-2 incorporated into the CPG layer of the eggshell rather than diffusing away (Fig. 5D), supporting the hypothesis that vitelline layer defects fail to retain soluble proteins in the perivitelline space between the eggshell and embryo. While the eggshells of PERM-2/4 depleted embryos retain some significant structural properties, the absence of these proteins results in a porous and improperly assembled eggshell.

## Discussion

The current work identifies three key proteins required for vitelline layer formation, to our knowledge the first such proteins to be described in nematodes. Before fertilization, CBD-1 recruits PERM-2 and PERM-4 to the nascent vitelline layer on the oocyte surface via two chitin-binding domains in its N-terminus. CBD-1 also anchors the EGG/CHS complex to the oocyte cortex through a domain distinct from the PERM-2/4 recruitment domain (Fig. 6A). After fertilization, CBD-1 maintains its interactions with PERM-2 and PERM-4 as the vitelline layer is remodeled into the outer layer of the eggshell (Fig. 6B), while its interactions with the EGG complex are relieved (possibly through competition with newly synthesized chitin polymer (Johnston and Dennis, 2011)), followed by the complex’s internalization and/or degradation (Maruyama et al., 2007; Parry et al., 2009; Stitzel et al., 2007). During vitelline layer formation and remodeling, PERM-2 and PERM-4 co-dependently maintain each other’s localization in order to establish a diffusion barrier and promote structural integrity of the eggshell.

**Figure 6.**
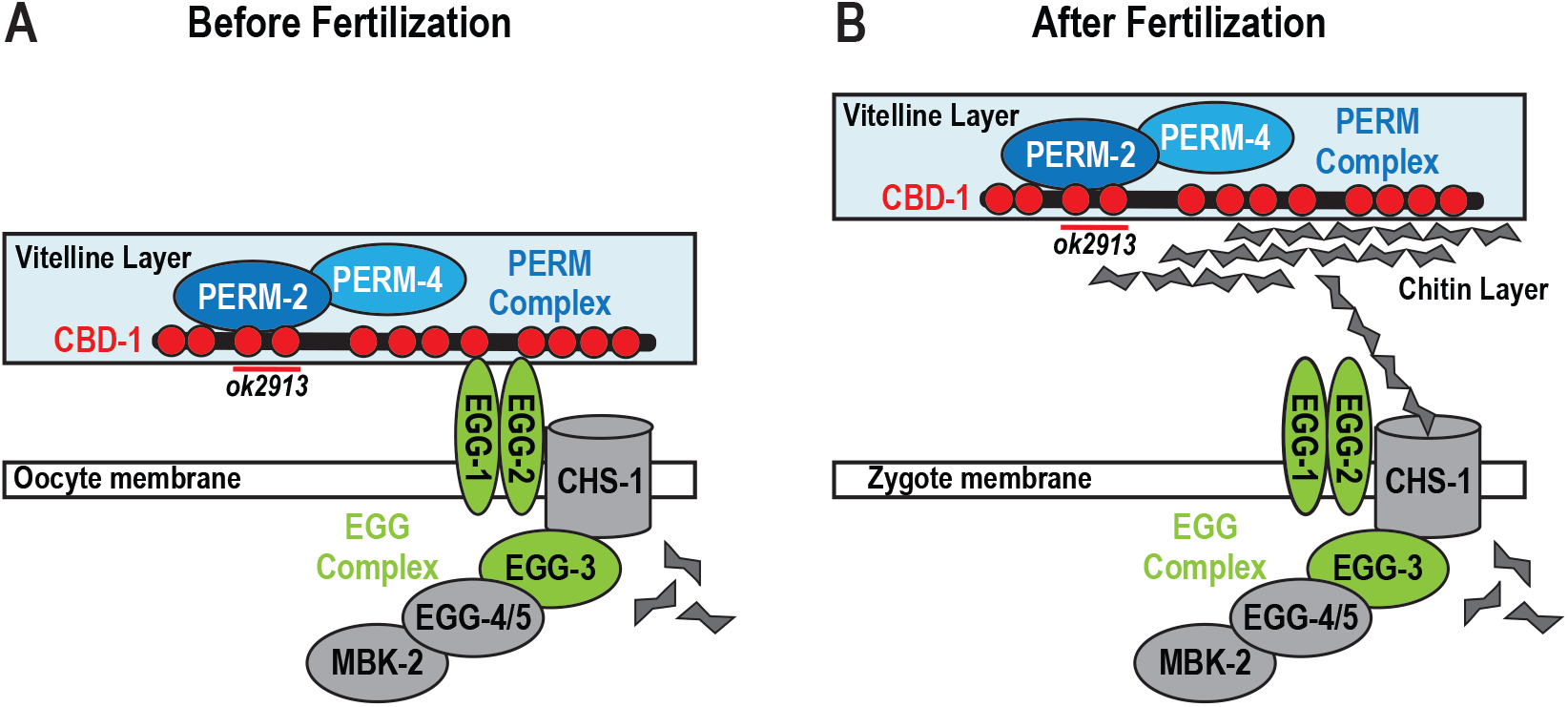
Model for vitelline layer remodeling in *C. elegans*. **(A)** Proposed structure of the vitelline layer before fertilization. CBD-1 (red) recruits PERM-2 and PERM-4 (blue) through two N-terminal chitin-binding domains (red circles, deleted by the ok2913 allele). CBD-1 also stabilizes the EGG complex (green and gray) through domains distinct from those required for vitelline layer organization. Members of the EGG complex are required for fertilization and egg activation. **(B)** After fertilization, synthesis of chitin may disrupt the interaction between CBD-1 and the EGG complex, allowing the vitelline layer to lift from the zygote surface. CBD-1 maintains association with the PERM complex in order to remodel the vitelline layer into a diffusion barrier that restricts entry and exit of large molecular weight compounds.

### CBD-1 is a multifaceted regulator of vitelline layer assembly

In the hierarchy of vitelline layer assembly, CBD-1 appears to function at the apex given (1) the severity of its phenotype compared to other components, and (2) its ability to recruit two functionally independent complexes via distinct domains. Embryos depleted of CBD-1 by RNAi produce a fragmented eggshell that slides off the embryo surface, exhibit polyspermic fertilization, and fail to undergo cell division (Johnston et al., 2010). By contrast, embryos co-depleted of PERM-2 and PERM-4 build rudimentary layers of the eggshell proper (vitelline, chitin, and CPG layers), and can complete several rounds of cell division before arresting around the gastrula stage, suggesting the PERM complex assembles and functions downstream of CBD-1.

As a large protein with 12 distinct chitin-binding domains, CBD-1 has the potential to interact with multiple factors on the oocyte surface to promote distinct aspects of early embryogenesis. In support of this idea, the *cbd-1(ok2913)* deletion allele that removes chitin-binding domains 3 and 4 near the N-terminus disrupts localization of the PERM complex, but not the EGG complex. By contrast, depletion of the entire CBD-1 protein by RNAi disrupts both the PERM and EGG complexes (Fig. 4, (Johnston et al., 2010)). Similarly, the phenotype of *cbd-1(ok2913)* mutants is less severe than that of *cbd-1(RNAi)* embryos, with the former capable of producing enough viable embryos to propagate the homozygous mutant strain (Fig. 1B), and the latter exhibiting polyspermic fertilization (Johnston et al., 2010) followed by complete sterility within 6 hours of RNAi treatment (data not shown). Given that two adjacent chitin-binding domains of CBD-1 can specifically promote PERM complex assembly, it will be important to explore the domains necessary to recruit EGG-1/2 and CHS-1/EGG-3 to the oocyte membrane in order to identify structure-function relationships important for nematode fertilization and egg activation.

### PERM-2 and PERM-4 promote structural integrity of the vitelline layer

The current model for *C. elegans* eggshell formation proposes that assembly occurs in a hierarchical fashion, where proper formation of outer layers is required for subsequent formation of inner layers (Olson et al., 2012). CBD-1 fits well with the model given that its inhibition prevents formation of the vitelline layer, chitin layer, CPG layer and permeability barrier ((Johnston et al., 2010) and Fig. 4CD). However, we were surprised to find that inhibition of the two other vitelline layer components, PERM-2 and PERM-4, had little effect on formation of the chitin and CPG layers (Fig. 5), and only inhibited downstream assembly of the permeability barrier (Fig. 1). These data suggest that CBD-1 is the main contributor to vitelline layer architecture and is central to promoting chitin layer formation after fertilization, either by regulating the distribution of chitin synthase on the oocyte membrane and/or organizing chitin filament assembly via its chitin-binding domains (Johnston and Dennis, 2011). By contrast, the role of PERM-2 and PERM-4 appears not to involve chitin layer deposition, but rather to promote formation of a diffusion barrier that can restrict the entry of extracellular factors and maintain secreted factors within the region between the embryo and eggshell that can serve important structural or signaling roles. For example, an externally-applied chitin probe stained the chitin layer more intensely in *perm-2(RNAi)* and *perm-4(RNAi)* embryos than in control embryos with an intact vitelline layer (Fig. 5A), suggesting that the vitelline layer is more porous when PERM-2/4 are absent. Similarly, the failure of *perm-2(RNAi)* and *perm-4(RNAi)* embryos to retain CPG-2 in the perivitelline space resulted in failure to properly assemble the permeability barrier (Fig. 5D), explaining how a defect in the outermost vitelline layer can impact the innermost permeability barrier of the eggshell, without affecting formation of the intervening chitin and CPG layers.

Inhibition of PERM-2/4 not only caused defects within the eggshell layers, but also led to abnormalities at the eggshell surface. Inhibition of PERM-2/4 caused abnormal adherence at the eggshell surface, such as when embryos dissected from depleted worms were extruded attached to one another (Fig. 5B). This string of embryos phenotype suggests that absence of PERM-2/4 either uncovers adhesive components on the eggshell surface, or alternatively prevents the binding of uterine secretions that are normally thought to adhere to the outer eggshell as another layer of protection (Stein and Golden, 2015; Zimmerman et al., 2015). The latter hypothesis is attractive, since it could also explain the enhanced staining of the chitin probe through its ability to more easily penetrate the external eggshell layers. It will be important to explore the vitelline layer surface at the ultrastructural level to determine the degree of remodeling following fertilization, and determine which aspects are dependent on PERM-2 and PERM-4 function. These types of studies can be compared to those in human embryos, where ultrastructural analysis shows a striking remodeling event that converts a porous zona pellucida into an inter-connected, cross-linked structure after fertilization (Magerkurth et al., 1999; Nikas et al., 1994).

### Mechanisms of vitelline layer remodeling include both evolutionarily conserved and species-specific features

In light of our findings on nematode vitelline layer assembly, we thought it interesting to consider which features are species-specific, and which share common principles across great evolutionary distance. Remodeling of the sea urchin vitelline layer into the fertilization envelope is one of the best understood and most visually dramatic examples in animals. Prior to fertilization, the sea urchin vitelline layer contains the egg bindin receptor proposed to interact with sperm, an extracellular isoform of the structural protein rendezvin, and the p160 protein that tethers the vitelline layer to the plasma membrane (Wessel and Wong, 2009). Fertilization triggers the exocytosis of cortical granules, which deliver structural building blocks and enzymes to aid in remodeling of the vitelline layer. For example, structural components including CUB- and LDLrA-domain proteins (e.g. proteoliasin, SFE1, SFE9, and additional rendezvin isoforms) associate in tandem arrays to expand the developing fertilization envelope, while the serine protease CGSP1 cleaves the p160 tether to release and lift the vitelline layer from the plasma membrane, aided by the hydration properties of sulfated glycosaminoglycans (Wessel and Wong, 2009). Finally, enzymes such as ovoperoxidase and transglutaminase crosslink structural proteins into a rigid fertilization envelope (Wong and Wessel, 2008b). Mining the *C. elegans* proteome for homologs of sea urchin fertilization envelope components failed to identify promising leads. While dozens of worm proteins contain CUB, LDLrA, and dual oxidase/peroxidase domains, none exhibit fertilization defects or eggshell permeability in genome-wide RNAi screens. Despite the lack of homology at the protein level, vitelline layer remodeling in worms and sea urchin do exhibit mechanistic similarities. For instance, in *C. elegans*, the vitelline layer is tethered to the embryo surface via interaction of the vitelline layer protein CBD-1 with the egg surface proteins EGG-1/2 and chitin synthase (CHS-1), similar to the p160 tether in sea urchin. In *C. elegans*, this tether is proposed to be reversed upon activation of chitin synthesis, with chitin competing for binding domains on CBD-1 to displace the EGG/CHS interaction (Johnston and Dennis, 2011). *C. elegans* requires an alternative mechanism to release the tether via substrate competition rather than protease delivery, since *C. elegans* cortical granules are not exocytosed upon fertilization, but rather 15 minutes later during anaphase of meiosis I. This is in contrast to sea urchin, where the CGSP1 protease is released from cortical granules immediately after fertilization, allowing rapid cleavage of the p160 tether.

The mammalian zona pellucida (ZP) is likewise analogous to the nematode vitelline layer. This thick extracellular coat resides on the surface of unfertilized oocytes and is remodeled following fertilization. The main structural components of the zona pellucida are ZP1-3 (ZP1-4 in humans), which contain conserved structural hallmarks called ZP domains (ZPD) (Wassarman and Litscher, 2016). In mammals, ZP proteins participate directly in sperm-egg binding (Bleil and Wassarman, 1980), which contrasts with *C. elegans* where depletion of CBD-1, PERM-2, or PERM-4 do not prevent fertilization. The other CBD-1 interactors, EGG-1 and EGG-2, are proposed to be involved in sperm-egg interaction and may fulfil this role (Kadandale et al., 2005), though additional sperm-egg recognition factors are still being sought. In addition to sperm recognition, vertebrate ZP proteins are involved in thickening the zona pellucida during oocyte maturation through polymerization of their ZPDs into cross-linked fibrillar structures (Darie et al., 2008; Jovine et al., 2002). Each ZPD contains ~260 amino acids, separated into ZP-N and ZP-C subdomains, with each subdomain stabilized by two disulfide bridges (Wassarman and Litscher, 2016). The *C. elegans* genome contains ~10 ZPD-containing proteins, but none are predicted to function during early embryonic development. While PERM-2, PERM-4 and CBD-1 do not share significant sequence homology with mammalian ZP proteins, it is interesting to note that each of the 12 CBD-1 chitin-binding domains is likewise stabilized by two disulfide bridges, reminiscent of the ZP-N and ZP-C domains. Likewise, two proteins of the sea urchin vitelline layer contain CUB domains that are stabilized through disulfide bonds (Wong and Wessel, 2008a), suggesting conservation of structural features across phyla. An interesting area of future research will be to investigate whether CBD-1, PERM-2 and PERM-4 form cross-linked fibrillar arrays to create a hardened eggshell, as is the case with the sea urchin CUB and LDLrA structural proteins and the mammalian ZPD proteins.

Our identification of the first set of vitelline layer components in the nematode eggshell provides an opening to explore both widely-conserved and species-specific strategies used by animals during the formation of protective extracellular barriers, and could help identify novel therapeutic targets to fight parasitic nematode infection through destruction of this highly impenetrable barrier essential for embryonic viability.

## Acknowledgements

The authors thank members of the Fall 2012 Pomona College Advanced Cell Biology class (Claire Brickson, Jessica Chiang, Vivian Chou, Frances Hundley, Amy Li, David Morgens, Tyler Oe and Brian Wysolmerski) for initial investigations with PERM-2 and PERM-4; David Levine and Andy Golden for worm strain AG212; Katrina Dank, Jacob Brawer and Adia Ja’Nea James for technical assistance; and Matthew Marcello, Karl Johnson and Fabien Jammes for thoughtful comments on the manuscript. Some nematode strains used in this work were provided by the Caenorhabditis Genetics Center, which is funded by the NIH Office of Research Infrastructure Programs (P40 OD010440).

## Author contributions

Conceptualization, SKO; Investigation, DPG, HVL, DP, ZTW, MCH, JAP, JKD, SKO; Writing – Original Draft, SKO; Writing – Reviewing & Editing, DPG, HVL, DP, ZTW, JJM and SKO; Supervision, SKO and JJM; Funding Acquisition, SKO and JRY.

## Abbreviations used in this paper

CBD: chitin-binding domain
CHS: chitin synthase
co-IP: co-immunoprecipitation
dsRNA: double-stranded RNA
EGG: egg sterile
MBK: mini-brain kinase
PERM: permeable eggshell
RNAi: RNA interference
SEC: self-excising cassette
SQV: squashed vulva
ZPD: zona pellucida domain

---

**Table S1.**
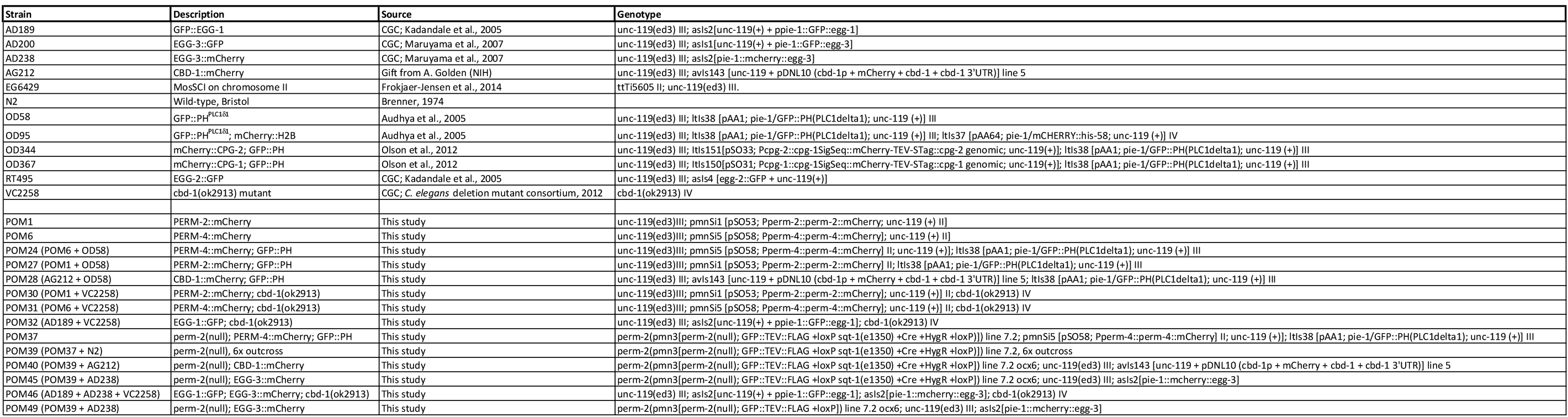
Worm strains used in this study.

**Table S2.**
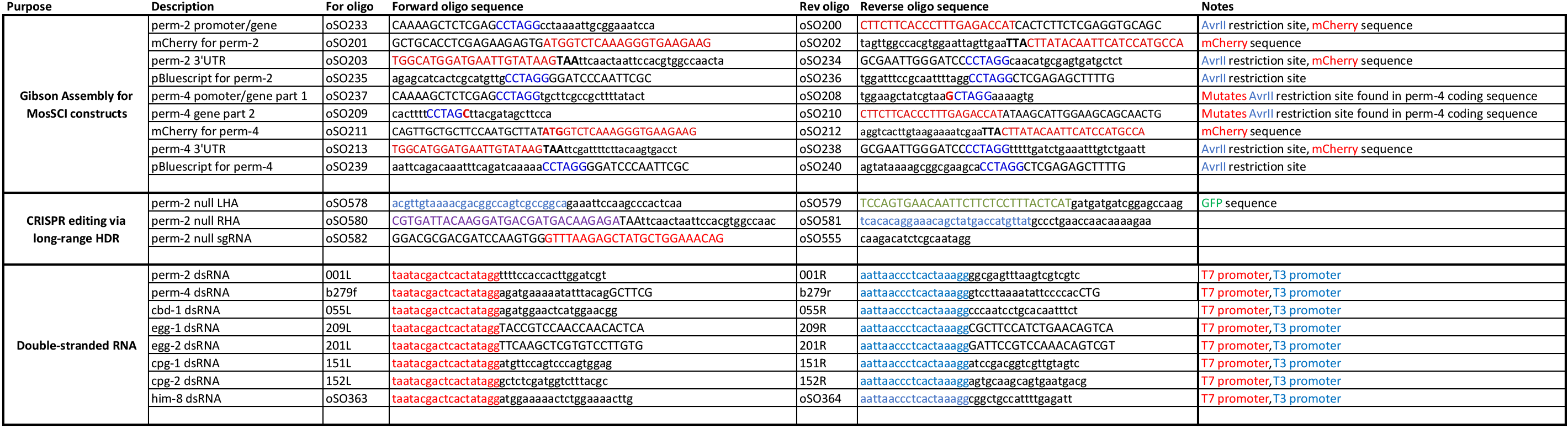
Oligos used in this study.

**Table S3.**
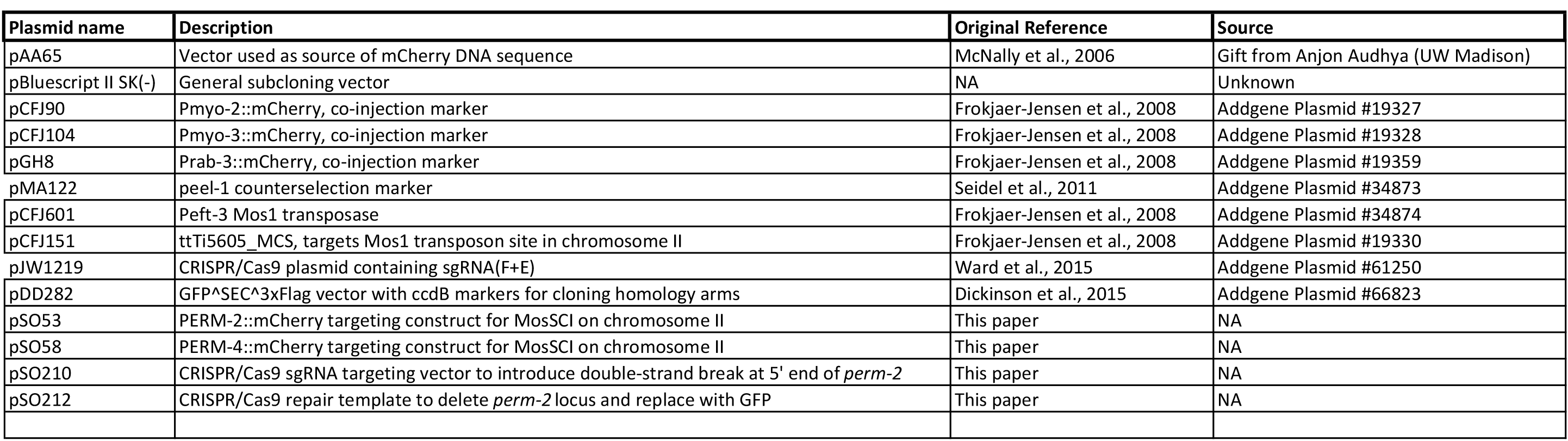
Plasmids generated and/or used in this study.

